# Feedback regulation by the RhoA-specific GEF ARHGEF17 regulates actomyosin network disassembly

**DOI:** 10.1101/2024.08.28.610052

**Authors:** Vasundhara Rao, Benjamin Grädel, Lucien Hinderling, Jakobus Van Unen, Olivier Pertz

## Abstract

We report that the RhoA-specific guanine nucleotide exchange factor ARHGEF17 localizes at the back of a fibroblast’s contractile lamella and regulates its disassembly. This localization emerges through retrograde ARHGEF17 transport together with actomyosin flow that most likely involves interactions with ATP-actin at F-actin barbed ends. During this process, ARHGEF17 increasingly oligomerizes into clusters that co-localize with myosin filaments, and correlate with their disassembly at lamella’s distal edge. ARHGEF17 loss of function leads to decreased RhoA activity at the lamella back and impairs its disassembly. High RhoA activity is however maintained at the lamella front where phosphorylated myosin light chain is observed. We propose that the low contractile, disassembling actomyosin network at the lamella back generates barbed ends leading to ATP-actin production, and ARHGEF17 binding. This then locally activates RhoA-dependent contractility, ensuring robust lamella disassembly through fracturing rather than re-inforcement. ARHGEF17 exemplifies the spatio-temporal complexity of Rho GTPase signaling and the requirement of feedback mechanism for homeostasis of contractile actomyosin networks.

## Introduction

Rho GTPases regulate cytoskeletal dynamics essential for cell/tissue morphogenesis (Etienne-Manneville and Hall, 2002). Their (in)activation are orchestrated by Guanine nucleotide exchange factors (GEFs) and GTPase Activating Proteins (GAPs). In their GTP-bound state, Rho GTPases activate downstream effectors to modulate cytoskeletal dynamics. Initial studies, based on population-averaged activity measurements and dominant mutants involved Rac1 in lamellipodial extension, Cdc42 in filopodial protrusion, and RhoA in myosin-driven contractility.

Fluorescence Resonance Energy Transfer (FRET)-based biosensors unveiled more complex regulation (Itoh et al., 2002; Kraynov et al., 2000; Nalbant et al., 2004; Pertz et al., 2006) in which RhoA/Rac1/Cdc42 exhibit specific spatio-temporal signaling patterns that fluctuate on time scales of tens of seconds during leading edge protrusion/retraction cycles (Machacek et al., 2009). Adding to this complexity, distinct RhoA/Rac1/Cdc42 signaling patterns are observed at the leading edge upon application of promigratory cues (Martin et al., 2016). The finding of such complex spatio-temporal Rho GTPase signaling patterns must involve precise regulation by GEFs/GAPs. Signaling pattern formation involves Rho GTPase flux, that consists of sequential spatially regulated events such as GEF-triggered Rho GTPase GTP-loading, its diffusion within the plasma membrane, and GAP-mediated inactivation. Ubiquitous GEF/GAP expression, observed across cellular systems (Rossman et al., 2005), is consistent with this complexity.

Comprehensive screening of their subcellular localization revealed that many GEFs/GAPs bind to cytoskeletal and adhesion structures (Müller et al., 2020), suggesting the existence of feedback regulation. By example, Rac1-specific GEFs versus GAPs respectively localize to adhesion structures at the front versus the back of the leading edge (Mueller et al., 2020). Thus, mechanical cues at adhesions at specific subcellular locations control the leading edge Rac activity gradient by spatially recruiting different GEFs/GAPs. Similarly, mechanosensitive interactions between the RhoGAP DLC1 and Talin allows its dissociation from focal adhesions under tension, locally increasing RhoA activity leading to positive feedback for robust and irreversible adhesion disassembly (Heydasch et al., 2023). Further, RhoGAP binding to F-actin provides spatial feedback shaping RhoA activity waves (Bement et al., 2024). Thus, interactions between GEFs/GAPs and cytoskeletal/adhesion structures provides feedback modalities that shape Rho GTPase signaling patterns. Traditional genetic manipulation techniques, which induce long-term changes in cellular states, do not adequately capture such spatio-temporal feedbacks. New methodologies are required to dynamically probe the Rho GTPase flux at adequate length and time scales in single cells.

Consistently with this signaling complexity, leading edge dynamics cycles involve two distinct yet interconnected structures, the lamellipodium and the lamella (Ponti et al., 2004; Small et al., 2002). The lamellipodium explores the environment, forming new protrusions and guiding the direction of movement. The lamella follows, consolidates protrusions into stable adhesions, and generates the contractile forces to pull the cell forward. This strongly suggests that multiple signaling modules must exist to spatio-temporally regulate these different cytoskeletal polymers.

Beyond Rho GTPase regulation, the self-organisation properties of the actomyosin cortex feed into regulation of morphogenetic processes. The actomyosin cortex can be understood as an active contractile matter that spontaneously form patterns such as waves, rings, or aggregates due to instabilities in the distribution of contractile forces (Salbreux et al., 2012). Experiments with isolated actomyosin systems have shown that the contractile activity of myosin motors can amplify small fluctuations in the distribution and alignment of actin filaments in a self-reinforcement manner that can result in structural changes such as buckling, fracturing or aggregate formation, that lead to tension loss (Ideses et al., 2018). Cells must therefore deal with emergence of potential catastrophic contractile instabilities in cells.

An interesting GEF that mediates crosstalk from the cytoskeleton to Rho GTPase signaling is the RhoA-specific GEF ARHGEF17, also known as Tumor Endothelial Marker 4 (Mitin et al., 2013, 2012). ARHGEF17 is characterized by an N-terminal actin-binding domain that exhibits a 90-fold higher affinity for ATP versus ADP-actin, and thus must bind to newly assembled F-actin filaments (Mitin et al., 2012; Prifti et al., 2022) (Fig. 1A). This is followed by a catalytic DH-PH domain, and a C-terminal WD40 propeller domain that might regulate protein-protein interactions. *In vitro* studies have shown that ARHGEF17 is specific for Rho isoforms (RhoA, RhoB, RhoC) over Rac1 and Cdc42 (Bagci et al., 2020; De Toledo et al., 2000; Mitin et al., 2012; Rümenapp et al., 2002a). This was later also found in a FRET biosensor assay that was used for mapping the specificities of all GEFs/GAPs for specific Rho GTPases in living cells (Müller et al., 2020). Expression of ARHGEF17 DH-PH domains leads to robust contractility (Mitin et al., 2012), further suggesting that it acts as a regulator of RhoA. ARHGEF17 therefore is highly likely to mediate cytoskeletal - Rho GTPase feedback, but the function of this feedback in the spatio-temporal regulation of signaling remains unclear.

**Figure 1.**
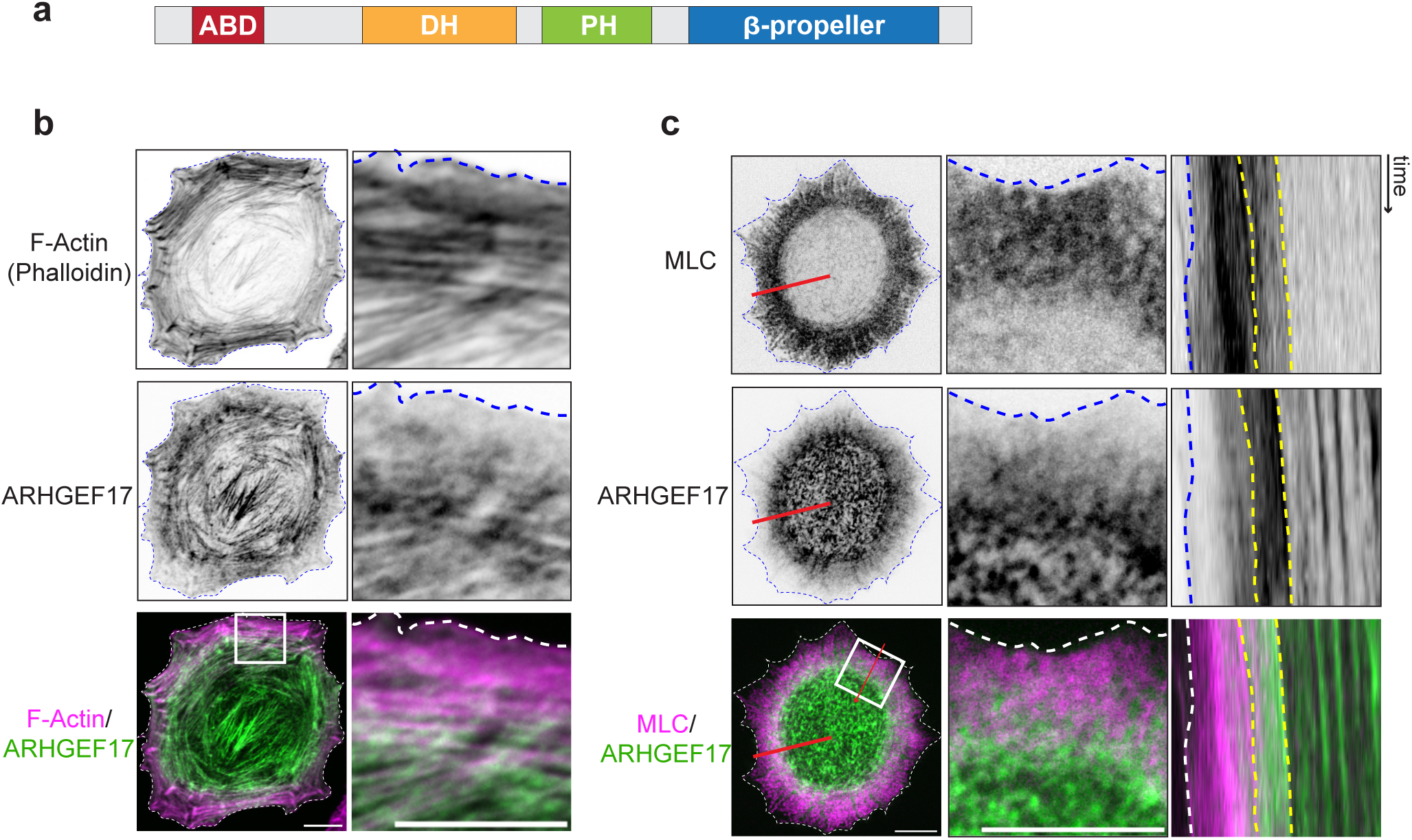
ARHGEF17 Structure, subcellular localization and dynamics. **A:** Scheme of the domain structure of ARHGEF17. (ABD : Actin Binding Domain) B,C,D: Images are shown with inverted black and white contrast (ibw), or overlayed channel color-coded images are shown. **B:** Confocal images of a spreading REF52 expressing ARHGEF17-mCherry and immunostained with Phalloidin. Left panels: whole cell. Right panels: magnified insets from boxes **C:** Spinning disk confocal timelapse imaging of a spreading REF52 cell expressing ARHGEF17-GFP, and MLC-mCherry. Right panel: Kymograph analysis of both signals along the line drawn in red in the left panel. Scale bars: 10 µm

Here, we show that ARHGEF17 dynamically localizes to the lamella back in spreading REF52 fibroblasts. ARHGEF17 oligomerizes from small to larger clusters as it travels with actomyosin retrograde flow. In the edge proximal lamella region, ARHGEF17 clusters sparsely bind to a highly contractile myosin network. In the edge distal lamella region, that is characterized by low contractility, ordered ARHGEF17 cluster arrays bind to transverse arcs, leading to disassembly of myosin filaments through their fracture. ARHGEF17 knockdown (KD) leads to lack of lamella disassembly, impeding symmetry breaking and polarization during spreading. On time scales of multiple hours, this leads to a very contractile stress fiber network. Optogenetic perturbation of contractility dynamics reveals that ARHGEF17 preferentially binds actomyosin networks in a relaxation state. As control cells, *ARHGEF17* KD cells still display a focused RhoA activity pattern at the edge, that has been shown to regulate lamellar contractility (Martin et al., 2016, 2014). However, decreased RhoA activity is observed at the back of this region, leading to lack of its disassembly. We propose that ARHGEF17 senses ATP-actin at barbed ends of actin filaments within the actomyosin lattice that is disassembling within the low contractile edge distal lamella. This locally activates RhoA-mediated actomyosin contractility that leads to fracture rather than reinforcement re-inforcement of the low contractile lamella, ensuring its efficient disassembly.

## Results

### ARHGEF17 binds to the edge distal region of the lamella during spreading

To study feedback from the cytoskeleton to Rho GTPases, we evaluated the subcellular localization of multiple GEFs and GAPs that were previously found to bind to F-actin or adhesion structures in REF52 fibroblasts (Müller et al., 2020). Confocal imaging of an exogenously-expressed GFP-tagged version of ARHGEF17 simultaneously together with F-actin staining, revealed that ARHGEF17 binds to the edge distal lamella region, as well as to low F-actin containing contractile structures in the middle of the cell (Figure 1B). Spinning disk confocal time-lapse imaging of ARHGEF17 and MLC dynamics during cell spreading again revealed a similar pattern (Figure 1C, Movie S1). Consistently with the fact that ARHGEF17 bears an actin binding domain (Mitin et al., 2012), it displayed retrograde flow together with MLC (Movie S1). Within the central domain of the cell, ARHGEF17 dynamically interacts with a contractile F-actin network with low MLC signal. Kymograph analysis revealed that the lamella exhibited two zones: a discrete band of strong MLC proximally to the edge, and a band of lower MLC signal distally to the edge that exhibits a sharp boundary where the transverse arcs disassemble (Figure 1C, separated by yellow dotted lines). ARHGEF17 displayed a strikingly different pattern: absence of signal in the lamellipodium with an increasing gradient that peaks in the low MLC edge proximal region, at the region where disassembly of transverse arcs occurs. These specific MLC and ARHGEF17 domains remain constant over time (Figure 1C). These data reveals a dynamic ultrastructure of the lamella with regions in which the myosin lattice adopts different densities, with ARHGEF17 prominently binding the edge distal region with low MLC density where lamella disassembly occurs.

### ARHGEF17 oligomerizes in cluster arrays that correlate with different actomyosin networks

To better understand ARHGEF17 subcellular localisation within the lamella, we imaged ARHGEF17 and MLC dynamics using structured illumination microscopy (SIM). SIM’s enhanced resolution revealed discrete ARHGEF17 punctate structures of different fluorescence intensities that we refer to as clusters, which might involve ARHGEF17 oligomerization through the C-terminal propeller domain (Mitin et al., 2012) (Figure 2A). Most likely through ARHGEF17’s actin binding domain, these clusters colocalized with actomyosin structures, and assembled into higher order arrays of clusters (Figure 2A-C). SIM resolved single myosin II filaments and their arrangement as stacks (Hu et al., 2017; Quintanilla et al., 2023) (Figure 2B, Movie S2). We then used an implementation of the Crocker-Grier centroid-finding algorithm (Crocker and Grier, 1996) to locate and quantify blobs that represent the myosin filament heads (Quintanilla et al., 2023) and the ARHGEF17 clusters, and measured distances between the latter (Figure S1A). Quantification of the distance between neighboring myosin filament heads, over the whole cell, revealed a distance of approximately 280 nm (Figure S1B), as previously observed (Hu et al., 2017; Quintanilla et al., 2024, 2023). ARHGEF17 clusters displayed a similar distance distribution (Figure S1B). The distance between neighboring myosin filament heads and ARHGEF17 clusters was 190 nm, suggesting that ARHGEF17 binding to F-actin positions them closely to the myosin filament heads (Figure S1B).

**Figure 2:**
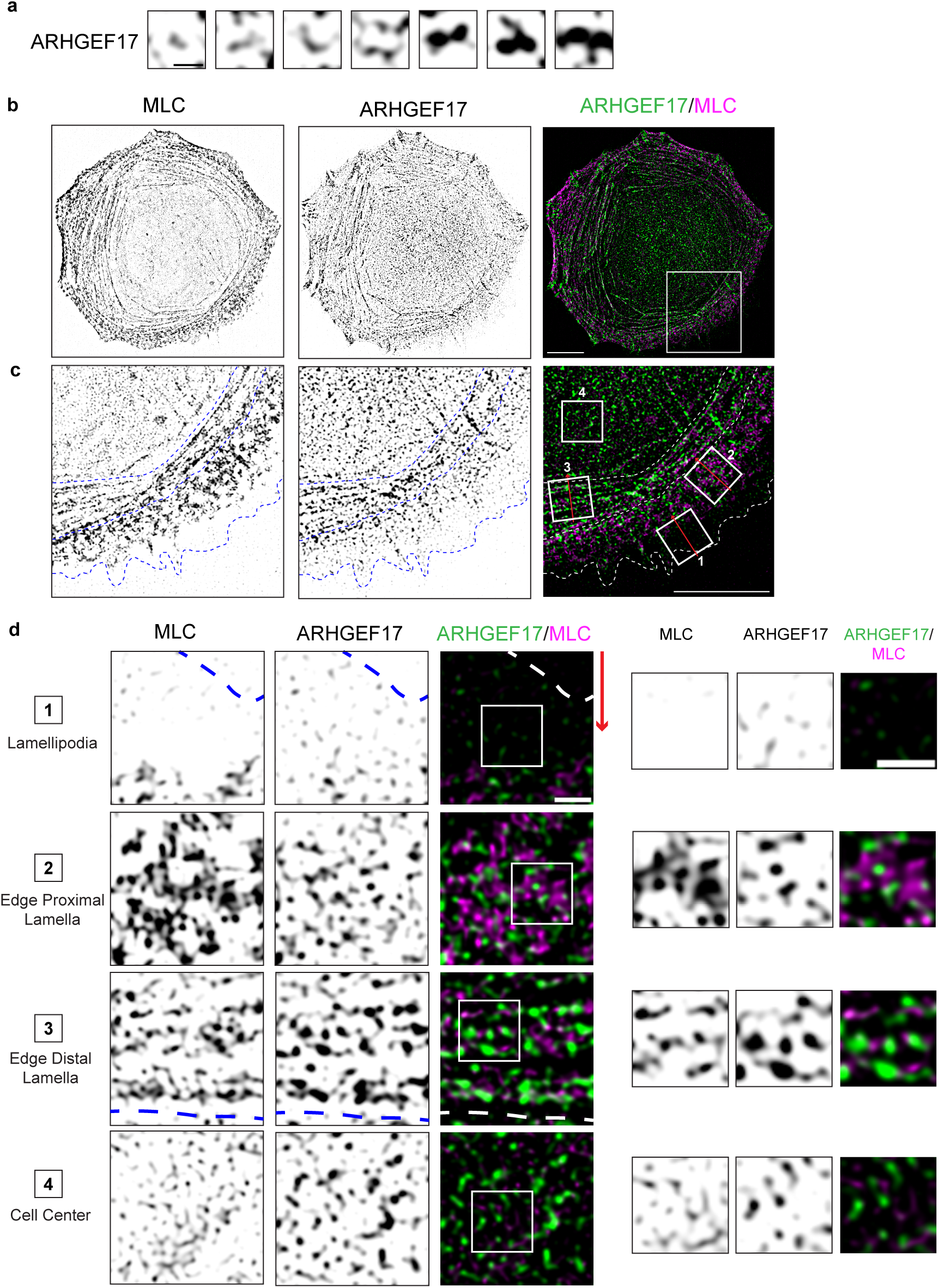
SIM imaging of ARHGEF17 and Myosin Light Chain subcellular localization. All images come from SIM imaging of a spreading REF52 fibroblast expressing ARHGEF17-GFP and MLC-mCherry. Ibw of single channels or color-coded overlays are shown. **A:** Gallery of ARHGEF17 clusters document their oligomerization properties. **B:** ARHGEF17/MLC subcellular localization over the whole cell. **C:** High magnification panel of A. Dotted lines delineate the cell edge, the interface between the contractile and low-contractile lamella regions, the low contractile lamella with the cell center. **D:** High magnification panels from C. Note that the panels have been oriented to be parallel to the edge, with the retrograde actin flow being oriented from top to bottom (denoted by the red arrow). Additional magnified insets on the right panels provide intuition about the ultrastructure of the MLC and ARHGEF17 arrays. Inset 1: Lamellipodium, inset 2: edge proximal contractile lamella, inset 3: edge distal low contractile lamella, inset 4: cell center. Scale bars: (A) 0.5 µm, (B,C) 10 µm, (D) 1 µm.

We then observed that the retrogradely flowing ARHGEF17 clusters displayed distinct organization patterns that correlated with different spatial arrangements of the actomyosin lattice. We first describe the different subcellular domains with specific actomyosin lattice arrangement. As expected, myosin II filaments were mostly absent from the lamellipodium (Figure 2B,C,D inset 1). In the edge proximal lamella region, a band of high density interwoven network of myosin filaments was observed (Figure 2D, inset 2). This crosslinked network most likely emerges from myosin redistribution to the branched lamellipodial actin that condenses into actin arcs (Burnette et al., 2011). Further within the cell, in the edge distal lamella region, we observed a second network of parallel myosin filaments with lower MLC density (Figure 2D inset 3). This explains the subdivision of the lamella in two zones with different MLC densities previously observed with confocal microscopy (Figure 1B,C). Confocal imaging of MLC and pMLC revealed that the edge proximal lamella contained a strong pMLC signal, while the edge distal lamella region contained much lower pMLC signals (Figure 3A,B,C inset 1). High resolution analysis of the edge distal lamella revealed that its parallel myosin arcs are decorated by sparse punctate pMLC puncta (Figure 3C inset 2). The cell center was mostly characterized by single myosin filaments that occasionally assemble into arcs, the latter retaining discrete pMLC punta (Figure 3C inset 3). Our blob quantification method revealed that the fluorescence intensities of the MLC signals were higher in the edge proximal versus distal lamella regions, and much lower in the cell center (Figure S1C-E). These results reveal that the lamella contains two separate myosin networks: 1. in the edge proximal lamella region, the myosin network adopts a highly interwoven structure and is under tension with high phosphorylated MLC, 2. the edge distal lamella region consists of a network of linear myosin filaments with mostly low tension. Here the myosin filaments’s motor domain might be trapped in a rigor state, and only act as a crosslinker without generation of contractility.

**Figure 3:**
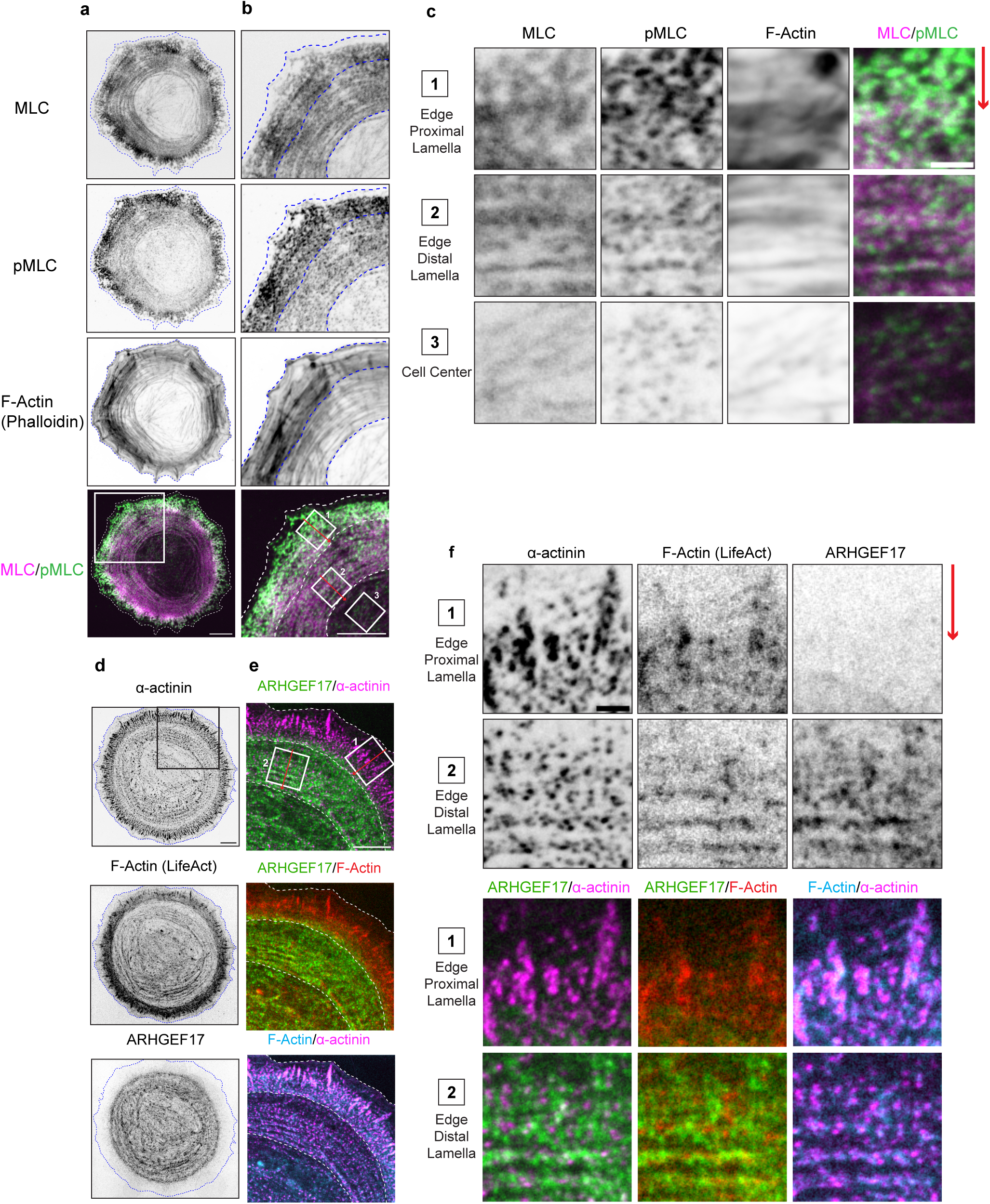
Subcellular localisation of MLC/MLC phosphorylation and ARHGEF17, F-actin and a a-actinin. A-C Confocal imaging of a spreading REF52 cell expressing MLC-mCherry and immunostained with pMLC and Phalloidin. **A:** Global view of the cell **B:** Magnification panel of the inset shown in (A) **C:** High magnification panels of the insets shown in (B). Inset 1: edge proximal contractile lamella, inset 2: edge distal low contractile lamella, inset 3: cell center. D-F: Confocal imaging of spreading REF52 cells expressing ARHGEF17-mCherry and LifeAct-miRFP and immunostained with a-actinin. **D:** Global view of the cell **E:** Magnification panel of the inset shown in (D) **F:** High magnification panels of the insets shown in (E). Inset 1: edge proximal contractile lamella, inset 2: edge distal low contractile lamella Ibw of single channels or color-coded overlays are shown. Note that the panels of (C) and (F) have been oriented to be parallel to the edge, with the retrograde actin flow being oriented from top to bottom (denoted by the red arrow). Scale bars: (A-B) 10 µm, (C) 2 µm, (D) 10 µm, (F) 2 µm

Evaluation of ARHGEF17 signals in these different subcellular regions revealed that ARHGEF17 forms arrays of clusters that exhibit increasing fluorescence intensities as they are transported retrogradely (Figure 2D). Small ARHGEF17 puncta were observed in the lamellipodium (Figure 2D, inset 1). In the edge proximal lamella region, ARHGEF17 assembled into larger clusters that localized in between the crosslinked myosin filaments (Figure 2D, inset 2). In the edge distal lamella region, ARHGEF17 clusters displayed a further increase in density and formed lattices that are parallel to the myosin filament network (Figure 2D, inset 3). ARHGEF17 clusters form networks suggesting that they bind multiple actin filaments simultaneously (Figure 2D, inset 3). Both myosin filaments and ARHGEF17 clusters defined a sharp boundary between the low contractile lamella region and the cell center (Figure 2C, dotted line). In the cell center, we observed homogeneous distribution of ARHGEF17 clusters of lower intensity than those observed in the edge distal lamella (Figure 2D, inset 4). Quantification of fluorescence intensities in detected blobs revealed that ARHGEF17 clusters in the contractile edge proximal have lower fluorescence intensity than those present in the low-contractile lamella region and the cell center (Figure S1C-E). We also evaluated the subcellular localization of ARHGEF17 with respect to F-actin and the actin crosslinker ⍺-actinin (Figure 3D-F) using confocal microscopy. In the contractile edge proximal lamella region, ⍺-actinin mostly bound to adhesions and diffuse ARHGEF17 signal were observed (Figure 3F inset 1). In the low contractile edge distal lamella region, we observed that ARHGEF17 clusters were colocalizing with F-actin filaments but were mutually exclusive with ⍺-actinin puncta where F-actin filaments are crosslinked (Figure 3F inset 2). This is consistent with the ability of ARHGEF17 to bind ATP-actin, mostly at the barbed-end (Mitin et al., 2012). We summarize the interactions of ARHGEF17 with the actomyosin lattice in Figure S2.

### ARHGEF17 clusters flow with the retrograde actin flow and correlate with disassembly of actomyosin transverse arcs in the lamella

Timelapse SIM imaging revealed that ARHGEF17 clusters flowed back with the actin retrograde flow (Figure 4A,B, Movie S2 and S3). During this process, we observed that the transported ARHGEF17 clusters exhibited fission and fusion events (Figure 4A,B, Movie S3), and increased in fluorescence intensity while they are transported back (Figure S1D,E). The correlation of ARHGEF17 transport by the actin retrograde flow, and the formation of ARHGEF17 clusters of increasing fluorescence intensities while transported centripetally strongly suggests a mechanism of advection that leads to coalescence of ARGEF17 into clusters. This flow was interrupted by disassembly of the myosin filament lattice in the lamella edge that faces the center of the cell (Figure 4C, Movie S4). An additional confocal timelapse dataset documents how ARHGEF17 enrichment correlates with transverse myosin arc disassembly (Figure S3, Movie S5). The high density of myosin arrays in the contractile edge proximal lamella region (Movie S3, upper panels) did not make it possible to quantify if a similar phenomenon occurs at this location. To further quantify the dynamic self organization properties of MLC and ARHGEF17, we evaluated the local correlation of the orientation of fluorescent signals using a custom algorithm inspired by the orientationJ plugin (Püspöki et al., 2016). This was performed for both channels independently. We observed that while the myosin lattice exhibited aligned arcs throughout the lamella, the characteristic aligned ARHGEF17 arrays were mostly observed in the edge distal lamella part at sites where transverse actomyosin arcs disassemble (Figure 4D, Movie S6). In the cell center, small ordered myosin arcs interacted with ARHGEF17 clusters which led to their immediate disassembly (Figure 4E). These datasets suggest the actomyosin retrograde flow dynamically organizes the advection and coalescence of ARHGEF17 clusters to concentrate the latter in transverse actomyosin arcs in the non-contractile lamella. This can also occur on actomyosin arcs transiently occurring in the cell center. When this spatially organizing mechanism of ARHGEF17 concentration exceeds a specific threshold, ARHGEF17 might activate the RhoA-Rho associated protein kinase (ROCK)-MLC activation to locally fracture the low contractile transverse arcs leading to their disassembly. Thus, local activation of RhoA through this spatially organizing mechanism might lead to an opposite effect than reinforcement of contractility which is usually the expected output of the Rho/ROCK pathway.

**Figure 4:**
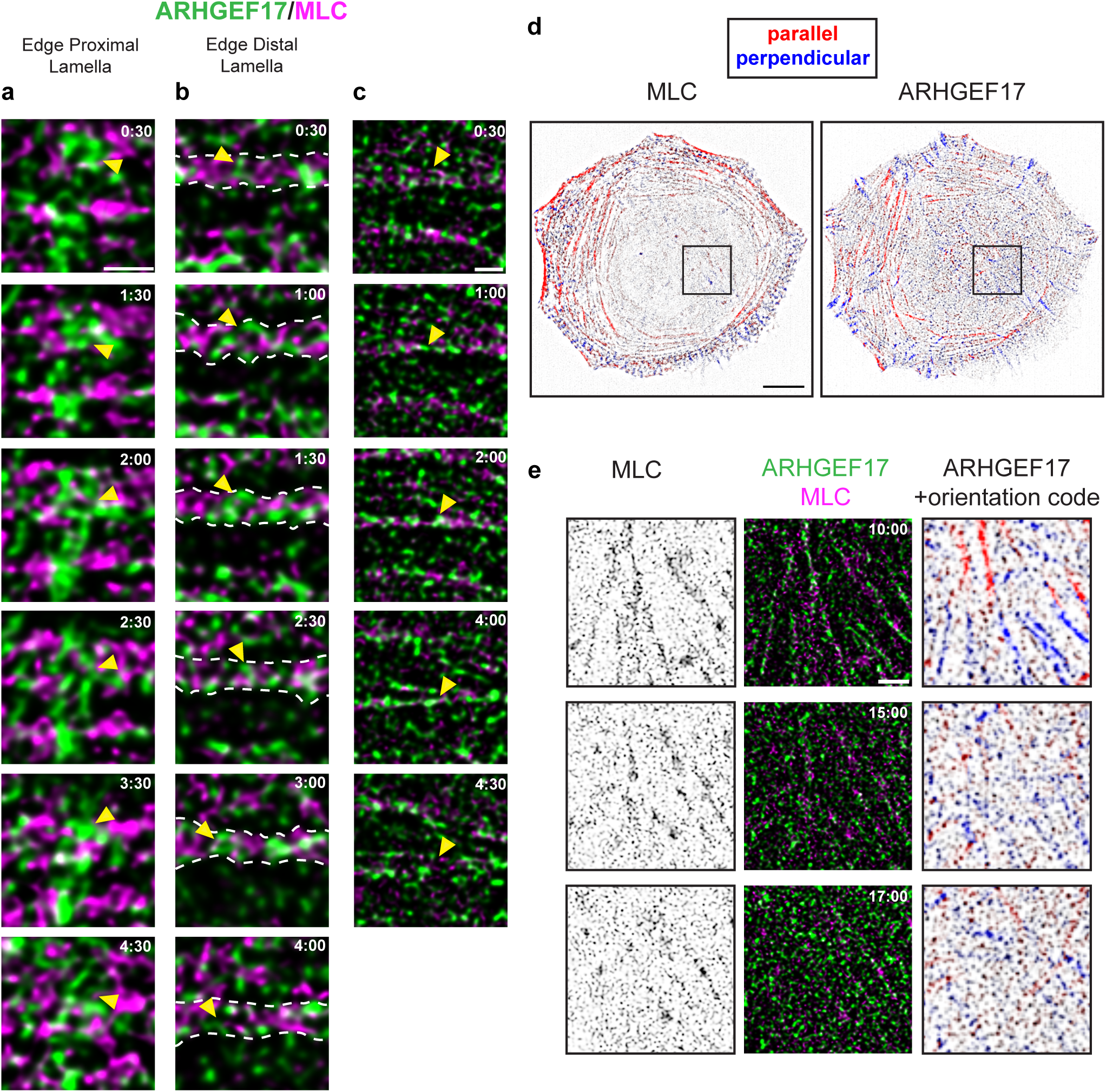
SIM imaging of ARHGEF17 and Myosin Light Chain dynamics. SIM imaging. SIM timeseries of the same cell shown in Figure 2 are shown. **A,B:** High resolution time series micrographs of ARHGEF17 and MLC dynamics in the edge proximal contractile (A) and edge distal low contractile lamella (B). **C:** Timeseries micrographs of ARHGEF17 and MLC dynamics in the low contractile edge distal lamella with an example of transverse arc disassembly. **D:** Analysis of correlated orientation of Myosin filaments (MLC) and ARHGEF17 clusters as analyzed using orientation-py. Correlated fluorescence signals that are parallel/perpendicular to the edge are color-coded in red and blue respectively. Ibw contrast is shown. **E:** Timeseries micrographs of the disassembly of an actomyosin fiber in the cell center by ARHGEF17. Left panel: MLC (Ibw contrats), Middle panel: ARGEF17/MLC overlay, right panel: ARHGEF17 + orientation code. Scale bars: (A-B-C) 1 µm, (D) 10 µm, (E) 2 µm

### ARHGEF17 loss of function leads to loss of actomyosin disassembly

To further evaluate ARHGEF17’s function, we performed RNA interference KD experiments, and evaluated F-actin, MLC phosphorylation and focal adhesion outputs during REF52 spreading (Figure 5A). We observed a KD efficiency of 88 % at mRNA level (Figure S4). To quantify these datasets, we built an image analysis pipeline to automatically segment cells and measure fluorescence intensity as a function of distance to the cell border (depicted in Figure S5A,B, see intensity profile analysis in methods section). During cell spreading, *ARHGEF17* KD cells displayed a prominent increase in F-actin signals in the edge distal lamella region (Figure 5A,B). This was accompanied by robust increase in focal adhesion size and intensity, which are hallmarks of high contractility. Surprisingly, this phenotype was only accompanied by a modest increase in phosphorylated pMLC given these robust contractile responses (Figure 5B). Time-lapse imaging of F-actin dynamics of *ARHGEF17* KD cells starting from the onset of cell spreading revealed that the latter remain spread longer than their control counterparts (Figure 5C, quantified in Figure 5D - area measurements, Movie S7). Further, *ARHGEF17* KD cells lacked the ability to break symmetry much longer than control cells, despite displaying a more contractile cytoskeleton (quantified in Figure 5D - solidity measurements). Approximately 12 hours after initial spreading, we observed that *ARHGEF17* KD cells display a very contractile actomyosin network with respect to their control counterparts (Figure 5C - 12 hours). To better understand the nature of the increased tension in the lamella cytoskeletal network in response to *ARHGEF17* KD, we imaged MLC using SIM. We observed increased stacking of myosin filaments in *ARHGEF17* KD cells in both the edge proximal and edge distal lamella, with the density of myosin filaments decreasing in the edge distal lamella region for both conditions (Figure 5E, quantified in Figure S5C). These results reveal that loss of function of the RhoA-specific GEF ARHGEF17, leads to loss of actomyosin disassembly in the edge distal non-contractile lamella. This leads to high contractility, as observed at the level of F-actin and focal adhesions, in presence of only a small increase in pMLC. After initial spreading, *ARHGEF17* KD cells display strong and exceptionally stable contractile arcs which might explain the lack of the ability to break symmetry for cell polarization. After multiple hours, *ARHGEF17* KD cells display an extremely contractile actomyosin network. SIM imaging reveals that *ARHGEF17* KD cells display increased stacking of myosin filaments. Together with the finding that ARHGEF17 recruitment to the actomyosin lattice leads to its disassembly (Figure 3), these results indicate that ARHGEF17 regulates disassembly of actomyosin networks in the non contractile edge distal lamella.

**Figure 5:**
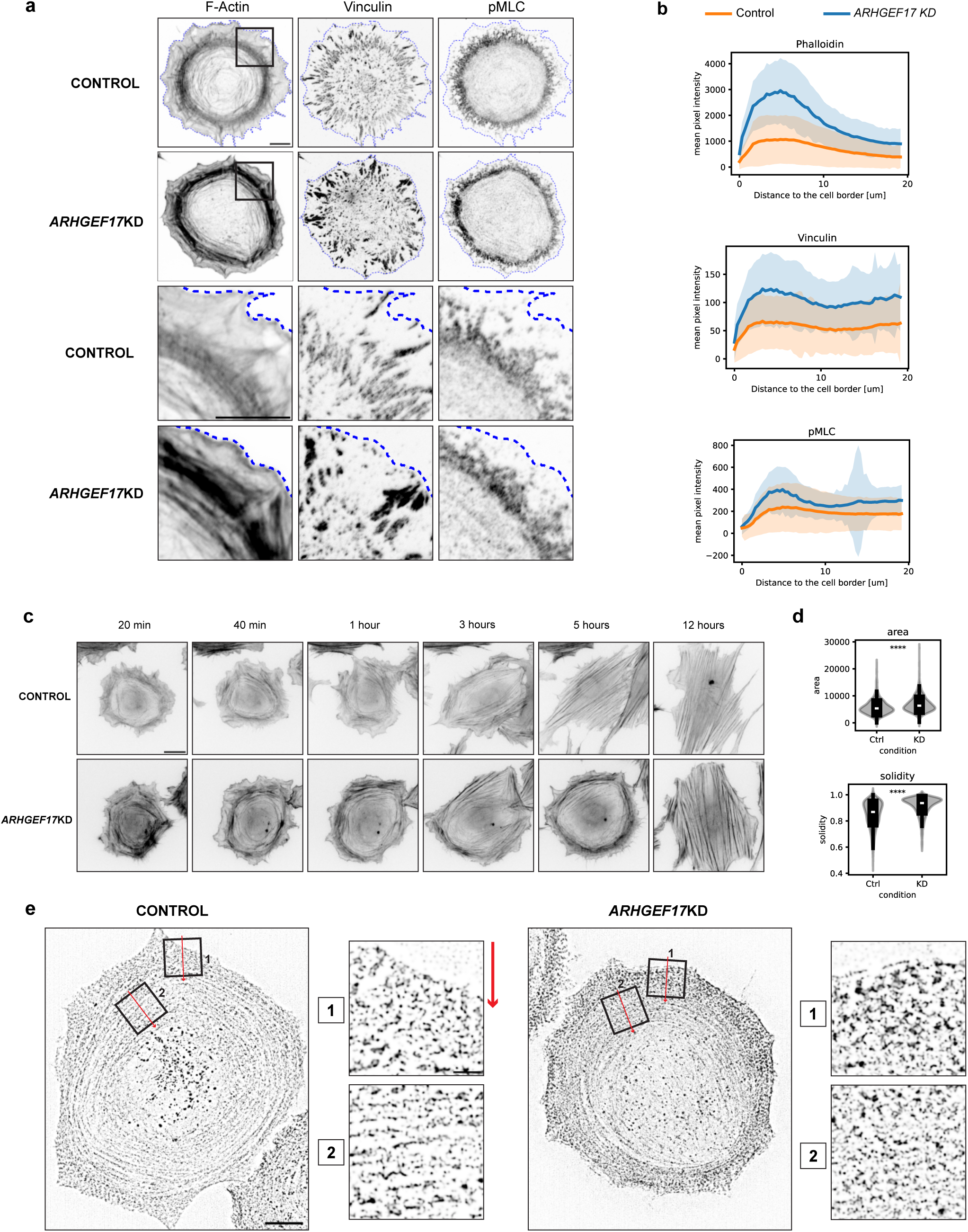
Effects of ARHGEF17 siRNA KD in REF52 cells. **A:** Fluorescence micrographs of cells stained with Phalloidin (F-actin), Vinculin and pMLC specific antibodies showing the impact of ARHGEF17 KD on cell morphology during spreading. Cells were detached, allowed to spread for 1 hour on fibronectin-coated coverslips. Images acquired using a 100x planApo oil immersion objective. **B:** Quantification of phalloidin, pMLC and vinculin fluorescence intensities in control and ARHGEF17 KD cells (Images acquired using a 100x objective). Distance intensity plots are shown for intensity of staining from cell border towards cell center. Shaded area indicates standard deviation. n= 477 cells (242 Control, 235 KD). **C:** Time-lapse series showing cell spreading and polarisation in control versus ARHGEF17 KD cells. Increased spreading and robust contractile F-actin structures are noted in KD cells within the first five hours as compared to control cells. At 12 hours, thicker contractile fibers are observed in ARHGEF17 KD cells. **D:** Quantitative analysis of area and solidity of control and KD cells after 5 hours of spreading, measured at time point 5 hours. Boxplot with interquartile ranges and superimposed violin plots are shown to provide an intuition about the measurement distributions. White bar indicates population average. n=2033 (945 Control, 1088 KD). Statistical test: Mann-Whitney U test. **E:** SIM images of spreading cells expressing MLC-mCherry in control and ARHGEF17 KD cells. High magnification panels showing (1) MLC in edge proximal lamella (2) MLC in edge distal lamella. The panels have been oriented to be parallel to the edge, with the retrograde actin flow being oriented from top to bottom (denoted by the red arrow). Scale bars : (A) 10 µm, (C) 5 µm, (E) 10 µm, (E) Insets: 2 µm

### ARHGEF17 loss of function decreases RhoA activity in the edge distal lamella

To evaluate how ARHGEF17 spatio-temporally regulates RhoA, we evaluated the effect of ARHGEF17 loss of function on RhoA activation patterns as measured by the FRET biosensor RhoA2G (Fritz et al., 2013). We simultaneously imaged F-actin dynamics using the lifeact construct. In spreading REF52 cells, we observed that as previously described (Martin et al., 2016, 2014), RhoA activity displays a focused region of RhoA activity directly at the leading edge, of similar size than the lamellipodium and the contractile edge proximal lamella as observed in Figure 3A-C (Figure 6, Movie S8). This was followed by a zone of lower RhoA activity (Figure 6A). As observed in figure 5A, *ARHGEF17* KD cells displayed an intense F-actin signal in the edge distal lamella, due to lack of their disassembly. In these cells, the edge localized RhoA activity pool remained unchanged, but lower RhoA activity correlated with lack of F-actin network disassembly in the edge distal lamella (particularly visible in the kymographs in figure 6B). Quantification by radially averaging RhoA activity across the periphery of the cell (Figure 6C,D) revealed that the RhoA activity pool at the edge remained unchanged, while the RhoA activity pool in the edge distal lamella is lower in perturbed versus control cells. Global RhoA activity levels did not change between control and perturbed cells, indicating that a very small pool of RhoA activity is affected by *ARHGEF17* KD (Figure 6E). The finding that both RhoA activity (Figure 6) and pMLC signals (Figure 5) remain present in the edge proximal lamella in *ARHGEF17* KD cells strongly suggests that this specific RhoA activity pool regulates lamellar contractility. The low contractile edge distal lamella might then emerge through retrograde flow in absence of RhoA/ROCK mediated MLC phosphorylation. The finding of lower RhoA activity in the edge distal lamella, that correlates with absence of F-actin network disassembly, indicates that ARHGEF17 regulates this specific RhoA activity pool. Thus, two spatio-temporal RhoA activity pools, one in the edge proximal and one in the edge distal lamella exist, and regulate actomyosin assembly and disassembly respectively.

**Figure 6:**
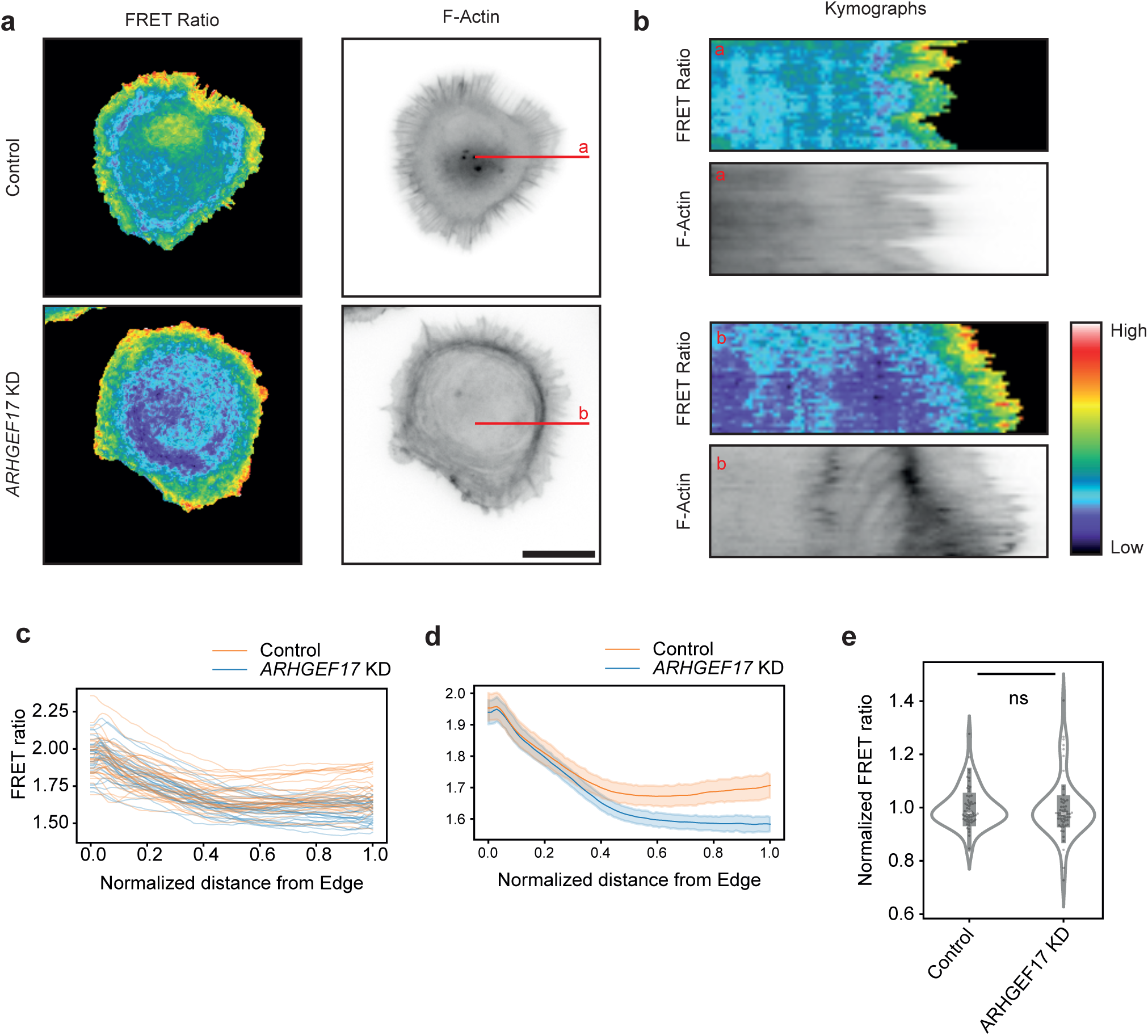
RhoA activity patterns in control versus ARHGEF17 KD cells. **A:** Ratio images of RhoA2G FRET signals and Lifeact-mCherry F-actin in control versus ARHGEF17 KD cells. Ratio images are color-coded according to the color scale shown in (B). Lifeact-mcherry image is shown in inverted black and white contrast. **B:** Kymograph analysis of the image in (A) along the red line. **C:** Distance intensity RhoA FRET ratio plots are shown for single cell (see material and methods image analysis section). n=34 cells per condition. **D:** Averaged distance RhoA FRET ratio plots. n=34 cells per condition. Shaded region indicates 95% confidence interval. **E:** Measurements of RhoA FRET ratio cell averages. Measurements are normalized to the median value of the control. A t-test was used to assess statistical significance. Scale Bar: (A) 50 µm.

### ARHGEF17 preferentially binds to relaxing actomyosin networks

The striking localisation of ARHGEF17 at myosin arcs, especially in the edge distal low contractile lamella, strongly suggests that ARHGEF17 binds to relaxed, disassembling myosin arcs (Figure 4). To test this hypothesis with an orthogonal approach, we used an optogenetic system to locally transiently activate Rho to evoke a temporary pulse of contractility followed by relaxation, and evaluate resulting ARHGEF17 dynamics. We used a re-engineered optoLARG actuator consisting of a nano SspB domain for efficient light-dependent recruitment of the LARG GEF domain to the iLID domain that was fused to a slowly diffusing stargazin membrane anchor (Heydasch et al., 2023) (Figure 7A). This construct was stably expressed at low levels together with ARHGEF17-mCherry and lifeact-miRFP F-actin (Figure 7A). Illumination of a spot with blue light led to local contractility as evidenced by accumulation of F-actin (Figure 7B, Movie S9). After blue light removal, contractility rapidly disappeared. Kymograph analysis revealed that ARHGEF17 was specifically binding to relaxing actomyosin networks. To further corroborate this finding, we imaged ARHGEF17 dynamics in response to myosin network relaxation triggered by Y-27632-mediated ROCK inhibition (Figure 7C, Movie S10). We found that after ROCK inhibition, ARHGEF17 was immediately recruited to the relaxing actomyosin network until it eventually completely disassembled. Together, these results strengthen the idea that ARHGEF17 selectively binds to actomyosin arrays that are in a state of relaxation.

**Figure 7:**
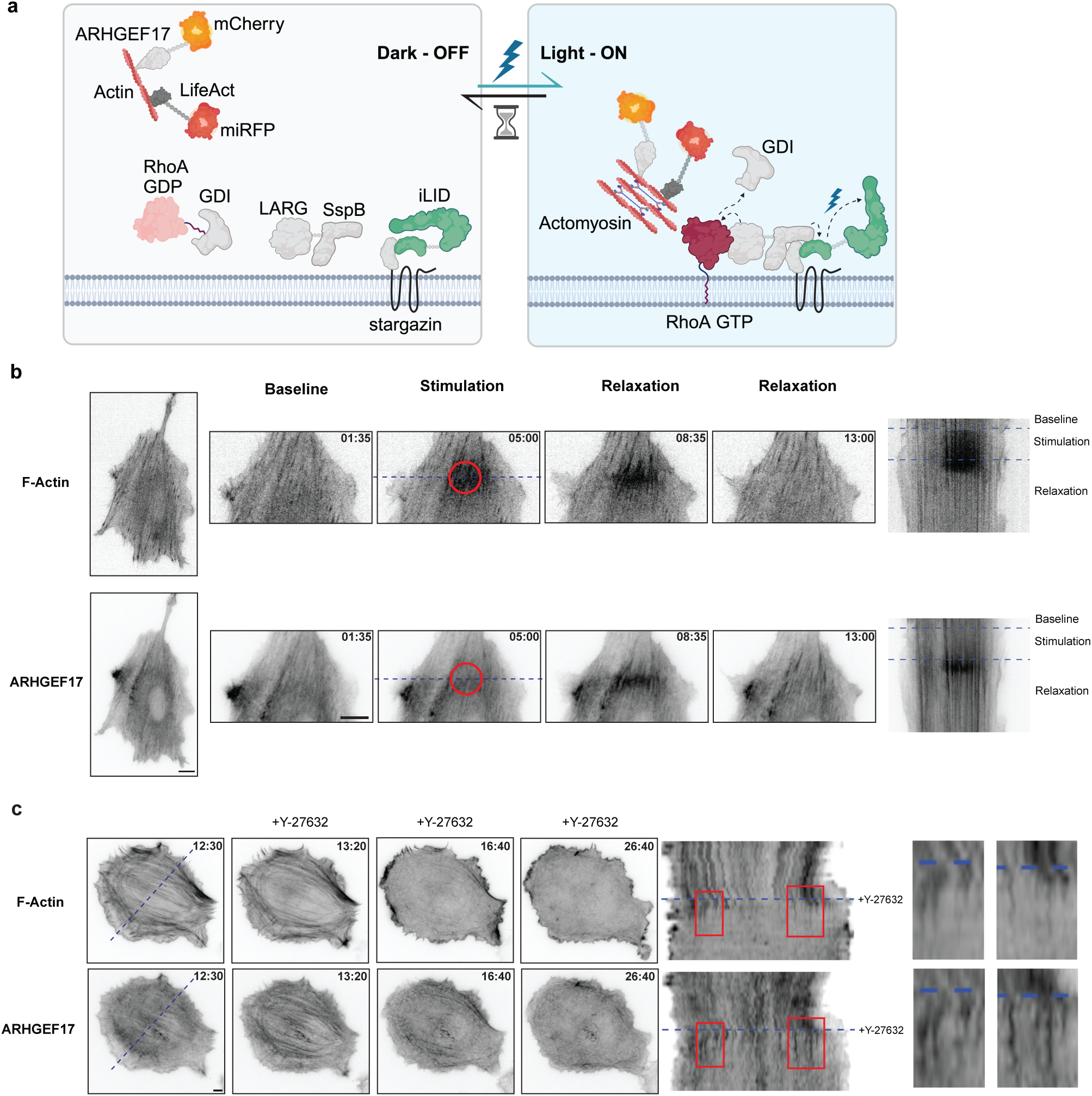
Mechanosensitivity of ARHGEF17 interaction with Actomyosin Network. **A:** Diagram of the optogenetic setup to study ARHGEF17 dynamics in response to a pulse of contractility. OptoLARG is an ILID-based optogenetic actuator to locally control Rho-mediated contractility and can be used with spectrally orthogonal mCherry tagged ARHGEF17 and the F-actin marker LifeAct-miRFP (adapted from Heydasch et al., 2023). **B:** Cells activated locally by blue light to induce actomyosin contraction and relaxation, illustrating ARHGEF17 binding during actomyosin disassembly rather than assembly. Time in mm:ss. **C:** Observations under ROCK kinase inhibitor Y-27632, showing ARHGEF17 transiently binds to disassembling actomyosin network. Time in mm:ss Scale bars: (B-C) 10 µm

## Discussion

While assembly of contractile actomyosin systems has been extensively studied, their disassembly remains virtually unexplored. It is currently assumed that cessation of Rho/ROCK signaling, which leads to activation of myosin light chain phosphatase (Amano et al., 2010) is sufficient for actomyosin network disassembly. Here, we report on a spatio-temporal signaling system dedicated to lamella disassembly. We show that ARHGEF17’s recruitment to the low contractile, edge distal region of the lamella, locally activates RhoA, and leads to its disassembly (Figure 8). This challenges the commonly accepted notion that RhoA solely regulates actomyosin network assembly, and shows that it can also control its disassembly. This underscores the need to understand Rho GTPase signaling in time and space and is consistent with the hypothesis of spatio-temporal signaling modules in which one GTPase can be spatially controlled by specific GEFs and GAPs to perform different functions in time and space (Pertz, 2010). A model on how ARHGEF17 regulates the disassembly of the lamellar actomyosin network is depicted in Figure 8.

**Figure 8:**
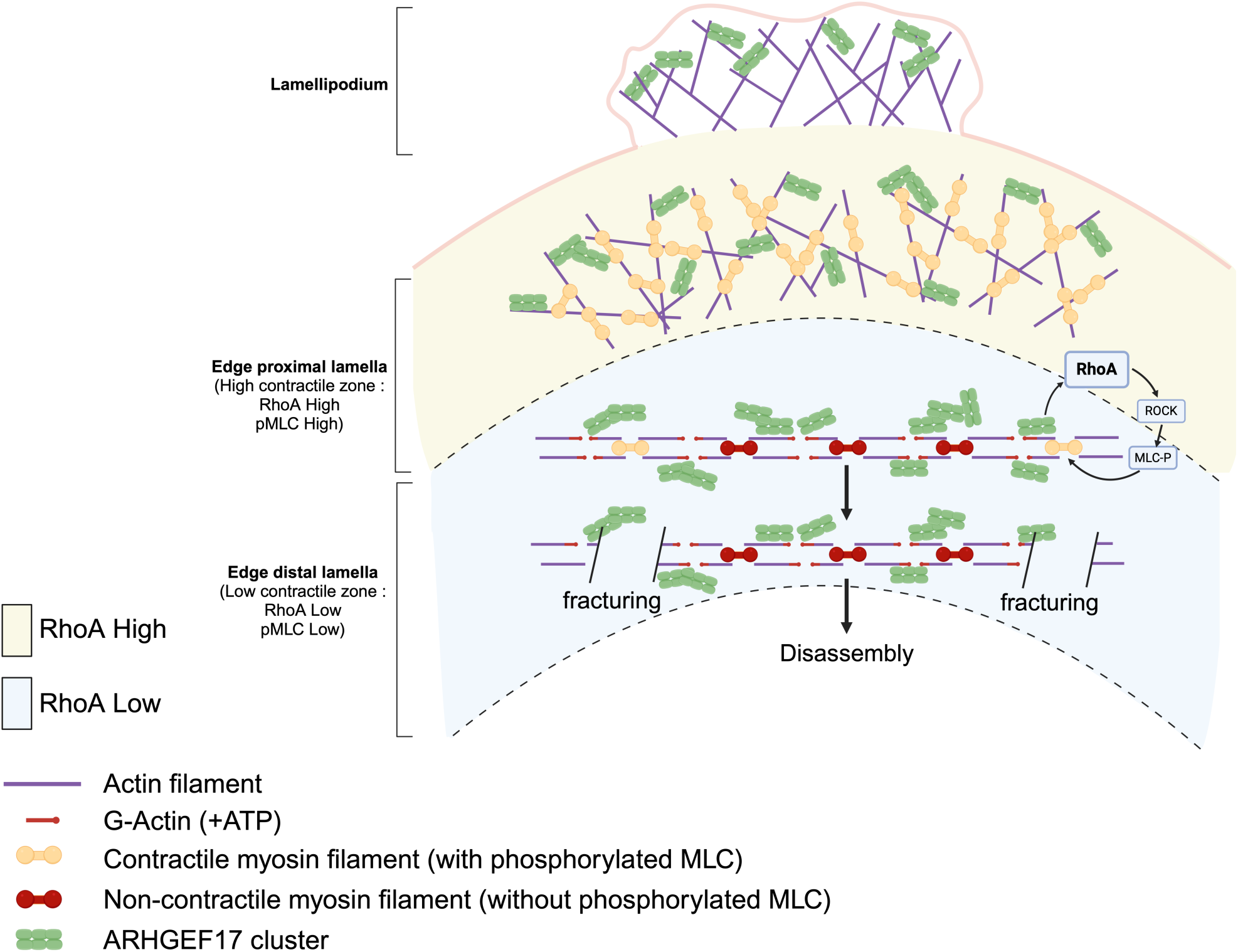
Model of ARHGEF17-mediated spatio-temporal regulation of lamella actomyosin network disassembly. Note that all myosin filaments and ARHGEF17 clusters are shown so as not to overload the model. 1. ARHGEF17 clusters start to interact with F-actin filaments in the lamellipodium and are transported backwards with the actomyosin retrograde flow. ARHGEF17 clusters most likely interaction with ATP-actin at barbed ends. 2. In the edge proximal lamella, RhoA activity regulated by GEFs/GAPs that remain to be discovered induce MLC phosphorylation through activation of ROCK, resulting in strong actomyosin contractility. ARHGEF17 clusters continue to be transported backwards, and increase in density through an advection mechanism. 3. In the edge distal lamella, the actomyosin network relaxes because it transitions into the region of lower RhoA activity, leading to MLC dephosphorylation. Most myosin filaments most likely only act as crosslinkers, without exerting tension. ARHGEF17 localizes to barbed ends of F-actin filaments crosslinked by both non contractile myosin filaments and alpha-actinin. High ARHGEF17 concentration locally applied to the non contractile actomyosin network leads to its disassembly, most likely by fracturing the actomyosin lattice.

An important component of this spatio-temporal signaling system is the mechanism by which ARHGEF17 is dynamically recruited to the lamella back region. SIM imaging of myosin networks and ARHGEF17 dynamics indicate that ARHGEF17 is transported backward with the actin retrograde flow (Figure 4A,B). This must involve the N-terminal domain of ARHGEF17 that shows high selectivity for ATP-actin, and thus barbed ends of F-actin filaments within the actomyosin lattice (Mitin et al., 2012) (Figure 8A). We observe that during this process ARHGEF17 clusters form through fusion-fission events (Figure 4A,B) and grow in size while they are transported centripetally (Figure 2, quantified in Figure S1C-E). This ultimately leads to highest ARHGEF17 density on linear transverse arcs in the edge distal lamella where the latter disassembles (Figures 2, 4C, S2, Movies S3-S4). Formation of ARHGEF17 clusters most likely involve the C-terminal β-propeller domain (Mitin et al., 2012), a domain regulating oligomerization in multiple proteins such as clathrin (Wood and Smith, 2021) or Cdc20 and Cdh1 that are involved in the activation of the anaphase-promoting complex/cyclosome during cell division (Peters, 2006). The observation that ARHGEF17 clusters bear similar size features than myosin filaments (Figure S1B), strongly suggest that these clusters contain tens of ARHGEF17 molecules. Our live imaging data suggests that ARHGEF17 clusters grow through an advection mechanism that involves interaction with F-actin filaments during retrograde flow. Note that ARHGEF17 also contains a PH domain that has been documented to bind to membranes (Mitin et al., 2012). Confining ARHGEF17 transport to the membrane might feed into this advection mechanism. An open question is why ARHGEF17 does not efficiently bind actin barbed ends in the lamellipodium. We speculate that interactions with F-actin together with the advection mechanism are necessary for concentration of ARHGEF17 clusters in the back lamella region. Once ARHGEF17 clusters disassemble together with the transverse myosin arcs in the distal lamella, they persist as disordered clusters in the cell center (Figure 2D, quantified in Figure S1C-E) where they control disassembly of spurious actomyosin networks in the cell center (Figure 4C).

Another important aspect of our spatio-temporal signaling system is how ARHGEF17 leads to actomyosin network disassembly during cell spreading. As just mentioned above, ARHGEF17 interactions allow its dynamic accumulation at the lamella back region where it correlates with actomyosin network disassembly (Figure 4). ARHGEF17 KD experiments (Figure 5) as well as FRET experiments indicated that ARHGEF17 regulates actomyosin disassembly through a RhoA-dependent mechanism (Figure 6). How does this workWe propose that local application of tension through ARHGEF17/RhoA/ROCK might fracture transverse arcs in the lamella back leading to their disassembly. This is consistent with *in vitro* (Murrell and Gardel, 2012) and *in vivo* (Wilson et al., 2010) evidence that myosin contractility can disassemble actin networks. This mechanism might be further influenced by the following factors: 1. Build-up of robust ARHGEF17 clusters might allow for a critical concentration of local ARHGEF17 to efficiently locally switch ON RhoA activity; 2. the position of ARHGEF17 clusters at the barbed end of F-actin filaments, away from ⍺-actinin mediated crosslinks which are crucial for stability of actomyosin networks (Dwyer et al., 2022), might be an advantageous weak point to fracture the transverse myosin arcs through local application of tension (Figure 8); 3. prominent ARHGEF17 clusters specifically localize within the low contractile edge distal lamella in which myosins might only act as crosslinkers, which might be easier to fracture (Figure 8B); 4. low contractile actomyosin networks in the edge distal lamella might already be prone to disassembly, producing new barbed ends for actin polymerization, ATP-actin incorporation, and thus further ARHGEF17 recruitment. Our optogenetic and drug perturbations of contractility (Figure 7) specifically illustrate the last point - during an optogenetically-evoked contractility transient, ARHGEF17 specifically binds to relaxing actomyosin networks (Figure 7A,B). Here, disassembling myosin networks might produce barbed ends that can incorporate ATP-actin, and recruit ARHGEF17 that provides a positive feedback by locally and transiently evoking RhoA-mediated contractility. An identical scenario applies to our ROCK inhibition studies (Figure 7C). Thus, the self-organization properties of ARHGEF17, that involve interaction with ATP-actin at F-actin barbed ends, as well as oligomerization might allow for its localization to low contractile, disassembling myosin networks at the lamella back region, providing a positive feedback for their robust disassembly.

Our data reveals the existence of two spatially distinct RhoA activation zones at the leading edge of the cell, the 1st regulating actomyosin assembly, and the 2nd regulating actomyosin disassembly. The 1st spatial pool of RhoA activity spans the lamellipodium and the contractile edge proximal lamella region, displays high pMLC signals (Figure 3A-C) and regulates actomyosin network assembly as already shown before (Martin et al., 2016, 2014). Rac1 activity is also present at that location (Martin et al., 2016), and RhoA activity might locally apply tension to lamellipodial branched actin network to switch on actomyosin retrograde flow. The finding that *ARHGEF17* KD maintains the 1st spatial pool of RhoA (Figure 6) indicates that another GEF and GAP must regulate it. Candidate RhoA-specific GEFs might be LARG and GEF-H1 that have been shown to regulate cytoskeletal responses in response to force application to integrins (Guilluy et al., 2011). The precise mechanisms that shape this focused RhoA activity pool remain to be discovered. The second ARHGEF17-mediated spatial RhoA activity pool is less intense and spans the non-contractile edge distal lamella region that displays sparse pMLC signals to regulate contractility for actomyosin network disassembly. Through retrograde flow, the contractile actomyosin network under tension in the 1st zone transitions in the 2nd zone of low RhoA activity, and adopts a relaxed state in which myosin filaments only displaying sparse pMLC signals, and thus most likely only acting as crosslinkers. This poorly disassembling actomyosin network can then activate ARHGEF17-mediated contractility through the mechanisms we have discussed above. Thus, there are two specific RhoA spatio-temporal activity pools that are regulating the lamella through different self-organization logics, with the ARHGEF17-regulated pool specifically dedicated for disassembly of actomyosin networks. Note that these RhoA activity patterns are particularly difficult to spatially resolve simply because active RhoA diffuses in the plasma membranes by virtue of its C-terminal geranylgeranyl moiety. Consequently, it is not possible to visualize ARHGEF17-mediated local MLC activation occurring at barbed ends in the low contractile edge-distal lamella. A better understanding of these different spatio-temporal RhoA activity pools will require single molecule imaging methods that have been previously used to image Rho GTPase dynamics (Mehidi et al., 2019).

Our work exemplifies the function of spatio-temporal feedback regulation in Rho GTPase signaling. Such feedback regulation is ubiquitous in Rho GTPase signaling, and for now has been mostly shown to be responsible for shaping spatio-temporal signaling patterns (Bement et al., 2024). In our work, the ARHGEF17 pathway can be understood as a spatially-regulated positive feedback to ensure robust and irreversible disassembly of actomyosin networks. This might be required for efficient homeostasis of the actomyosin cytoskeleton warranting the absence of contractile instabilities that might lead to actomyosin network buckling, fracturing, or the formation of aggregates as observed in cell free systems (Ideses et al., 2018). Further illustrating the concept that Rho GTPase feedback mechanisms are important for cellular homeostasis, we have recently observed a similar positive feedback mechanism impinging on RhoA activity that involves spatio-temporal regulation of the RhoGAP DLC1 at focal adhesions (Heydasch et al., 2023). Here, the positive feedback emerges from a mechanical input at focal adhesions, which locally amplifies RhoA signaling, and is required for irreversible focal adhesion disassembly. The finding that 30% of Rho GEFs/GAPs bind to the cytoskeleton or adhesions (Müller et al., 2020) strongly suggests widespread mechanisms of cytoskeletal to Rho GTPase feedback that might regulate cytoskeletal homeostasis or Rho GTPase pattern formation (Bement et al., 2024). Future elucidation of the role of GEFs and GAPs in spatio-temporal Rho GTPase signaling will need to take into account the concept of feedback regulation.

## ONLINE METHODS

### Cell Culture and Transfection

Rat Embryo Fibroblast 52 (REF52) cells were cultured in Dubelcco’s Modified Eagle Medium with 4.5 g/L glucose, 10% 4mM L-Glutamine, and 100 U/mL penicillin/streptomycin. Cells were grown at 37°C and 5% CO2.

For transfection, REF52 cells (45000 per well) were seeded in a 6-well plate a day before. Cells were transfection using Lipofectamine 3000 Reagent and following the protocol (Thermo Fisher). Transfected cells were kept overnight and visualised the next day.

### siRNA KD

4 ul of Lipofectamine RNAiMax (Thermo) was added to 500 ul of Opti-MEM (Thermo) and siPOOL solution for 3 nM final siRNA concentration. This mix was added to a well of 6-well plate and reverse transfection was done by adding 120000 cells in 1.5 mL medium volume in the well. After 24 hours, cells were re-seeded on glass bottom well plates (Celvis) which were coated with 5 μg/ml of human plasma fibronectin purified protein.

### DNA constructs

The optoLARG construct used was: pB3.0-optoLARG-mVenus-SspB-p2A-stargazin-mtq2-iLID, as described in Heydasch et al., 2023. ARHGEF17-mCherry construct was provided by Olivier Rocks, and ARHGEF17-GFP construct was from addgene #58893. MLC-mCherry, MLC-miRFP, LifeAct-miRFP and LifeAct-mNeonGreen were prepared previously in the lab.

### Optogenetic Experiments

REF-52 cells with optoLARG construct and LifeAct-miRFP were transfected with ARHGEF17-mCherry (kindly provided by Olivier Rocks) and plated on glass bottom well plates (Celvis) which were coated with 5 μg/mL of human plasma fibronectin purified protein (Merck) for one hour at room temperature. Cells were seeded at 6000 cells/well into 12 well plates and allowed to attach for 12 hours. Before imaging, FluoroBrite DMEM medium was added to the cells. NIS-Elements (Nikon) was used on the microscope for observing ARHGEF17 and LifeAct dynamics. Optogenetic stimulation was localized to specific regions using an Andor mosaic 3 DMD and the ROIs were individually defined for the cells. The images were taken using a 60x oil objective. mCherry-ARHGEF17 and miRFP-LifeAct were imaged for 20 cycles with 5 seconds time interval for baseline, then stimulation with blue light in the ROI for 65 cycles with 5 seconds time interval, followed by no stimulation acquisition with 5 seconds time interval.

### Immunofluorescence antibody staining

Vinculin, pMLC, Phalloidin-Atto647N (1:200 dilution), Alexa-Fluor-conjugated secondary antibodies (used as 1:1000) were from Molecular Probes. One hour after seeding, cells were fixed with 0.2% cold paraformaldehyde for 10 min, washed with 1x PBS three times, permeabilized with 0.1% Triton X solution, washed again with 1x PBS, and blocked for 30 min at RT with 2% BSA solution in PBS. Cells were incubated overnight with the primary antibody, and then washed with 2% BSA solution in PBS three times next day. Secondary antibody incubation was done for 1 hour at RT followed by three washes with 1x PBS.

### Y-27632 washout experiment

REF-52 cells with LifeAct-miRFP were transfected with ARHGEF17-mCherry and seeded on a 24-well glass bottom plate coated with 5 μg/mL fibronectin. Cells were allowed to attach for 12 hours and FluoroBrite DMEM medium was added later for imaging. Live cell imaging was done for addition of Y-27632. mCherry-ARHGEF17 and miRFP-LifeAct were imaged for 20 cycles with 5 seconds time interval for baseline and then 32 µm of Y-27632 was added. Imaging was continued with 5 seconds time interval until complete disassembly of actomyosin.

### RNA Extraction

Cells growing in a 6-well plate were rinsed with PBS and 1 mL TRI-Reagent per well was added. After 3 minutes of incubation, cells were transferred to a 15 mL tube and 200 µL chloroform was added. The tube was vortexed for 30 seconds and kept standing for 5 min at RT. After this it was centrifuged at maximum speed for 5 min in a centrifuge pre-cooled to 4°C. For each sample, 2 µL glycogen (20 mg/mL) was pipetted in fresh eppendorf tubes. From the centrifuged samples, 350 µL of upper aqueous phase was carefully removed and transferred into the corresponding tube containing glycogen. 600 µL of isopropanol was then added to each tube and mixed by inversion. The tubes were kept for 10 min at RT and then centrifuged at 4°C at maximal speed for 15 min. The supernatant was carefully removed and discarded. The pellet was gently rinsed with 1 mL ethanol 80% (diluted with DEPC water). The tube was centrifuged for 5 min at maximal speed and supernatant was discarded. The pellet was left to dry for 15 min and then 40 µL DEPC water was used to dissolve the pellet. The extracted RNA was kept on ice and concentration measured in a Nanodrop spectrophotometer.

### Reverse Transcription and qPCR

To reverse transcribe all polyadenylated mRNAs of a cell an oligo(dT) primer can be used which hybridises to the poly(A) tails of the mRNAs. For primer annealing, for each reaction, 5μg RNA + 1μl oligo (dT) primer 10μM were mixed and filled up to total 25 μl with water. This was incubated for 5 min at 75°C and then let stand at RT for 5 min. 25 μl of 2X reverse transcription complementation mix was prepared for each sample. For 25 μl 2X RT mix, 10.9 μl H2O, 5 μl 10X Affinity-ScriptTM buffer, 5 μl 100 mM DTT, 2.5 μl dNTP Mi× 10 mM each from NEB, 0.8 μl RNAse inhibitor (Dundee Cell Products) and 0.8 μl Reverse Transcriptase (Affinity Script Kit, Agilent/Stratagene) were mixed. The 25 μl 2X RT mix was added to each sample with RNA and annealed oligo(dT) primer, total 50 μl per tube. This was incubated for 10 min at 25°C (primer extension), 40 min at 42°C (reverse transcription) and 20 min at 60°C (enzyme inactivation). The resulting cDNA can be stored at −20°C. For qPCR, following primer pairs were used. GAPDH Fwd: GTCTCCTCTGACTTCAACAGCG, GAPDH Rv: ACCACCCTGTTGCTGTAGCCAA, ARHGEF17 Fwd: CATAGCTTCTCCCAACCGCA, ARHGEF17 Rv: AGGTACGCAGGAAAGCTCAC.

For 20 μl reaction, 10 μl of SYBR™ Green PCR Master Mix 1X (Thermo Fisher), 0.4 μl each of 10 mM Fwd and Rv primers, 15 ng of cDNA and water upto 20 μl were mixed and added to qPCR plate. Following program was run: Enzyme activation at 95 °C 1 min (1 cycle), Denature at 95 °C during 3 sec, Anneal/Extend/Acquire at 60 °C 15 sec. 40 cycles were done and data analysed.

### Structured illumination microscopy (SIM)

8-well chambered slides (Ibidi) coated with 5 μg/mL fibronectin were used for live cell imaging using SIM. REF-52 cells with MLC-mCherry were transfected with ARHGEF17-GFP (addgene #58893) and seeded in 8-well chambered slide. 30 min after seeding, FluoroBrite DMEM medium was added and cells were imaged using 63x oil objective using 488 and 561 laser. The interval time between images is 30 seconds. The processing of the acquired data was done using Zen 3.0 imaging software. The SIM Imaging was done at the Center for Microscopy and Image Analysis at the University of Zürich.

### RhoA activity FRET imaging

8-well chambered slides were coated with 5 μg/mL fibronectin. REF52 cells were seeded in Fluorobrite DMEM supplemented with 0.5% FBS, 0.5% BSA, L-glutamine and P/S. After 1 h images were acquired on a Nikon Eclipse Ti inverted microscope with a 60x Plan Apochromat Objective, using a Prim95B sCMOS camera with 2×2 pixel binning.

Donor and FRET channels were excited using a Lumencor Spectra X 440 nm LED and imaged sequentially with excitation filters 430/24 and a Dichroic Q465 long-pass filter. For donor emission, a 480/40 nm bandpass filter was used, and for FRET emission, a 535/30 nm bandpass filter was used.

Data was analyzed with custom python code with the pipeline described in (Spiering et al., 2013).

### Image analysis

Standard Python libraries were used for data processing and visualization (NumPy 1.24.4, pandas 2.0.3, skimage 0.22.0, Matplotlib 3.8.2).

#### SIM cluster analysis (Figure S1 and S4C)

For the analysis of clusters in SIM images in Figures 2 and 5, blob-like structures were detected using the Crocker-Grier centroid-finding algorithm (Crocker and Grier, 1996) implemented in the Python library Trackpy (version 0.6.1, https://zenodo.org/records/7670439). The expected diameter was set to 3 pixels, and the image was inverted to match the input format expected by Trackpy. For the distance analysis, nearest neighbors were detected by constructing and parsing a KD tree, using the algorithm described in (Maneewongvatana and Mount, 1999), which is implemented in the Python library SciPy version 1.9.1 (Virtanen et al., 2020).

#### Linear structure enhancement and orientation analysis (Figure 4D,E)

Linear structures and their orientations were detected by convolving the image with a 2D Gabor filter bank (Granlund, 1978). For all pixels, the vector pointing towards a manually placed cell centroid was calculated, and the orientation was derived as an angle from this vector. This angle was then compared to the angle of the Gabor filter with the highest response. Aligned angles (more pointing towards the cell center) were colored in blue, while non-aligned angles (more parallel to the cell edge) were colored in red. Final visualization was done using an HSI (hue, saturation, intensity) model inspired by the software OrientationJ (Püspöki et al., 2016). The blue/red values were used as hue, the Gabor filter response as saturation, and the original image as intensity values. The code to reproduce the figure is available on github as a Jupyter notebook (https://github.com/hinderling/orientation-py).

#### Intensity profile analysis (Figure 5B)

Cells were segmented with Cellpose (Stringer and Pachitariu, 2024) using the cyto3 model, a diameter of 150 px, and Phalloidin and DAPI as input channels. Cells touching the image border were excluded from the analysis. A Euclidean distance transform was applied to the cell masks to find the distance from the cell border for each pixel. For each cell, pixels with the same distance were binned, and the mean intensity was calculated for all background-subtracted fluorescence channels. The first 60 pixels (∼20 μm) were chosen for plotting. Cells with a shorter maximal distance were excluded from further analysis (mean maximal distance for all cells was 69.56 px).

### Statistics

Statistics were computed with scipy (version 1.9.1).

## Author contributions

VR and OP conceptualized the paper, VR and BG performed experiments, BG and LH performed data analyses and wrote code, JVU built the optogenetic toolkit, VR and OP wrote the paper.

## Acknowledgements.

We thank Daniel Riveline, Karsten Kruse, Damian Brunner and Miguel Vicente-Manzanares for constructive comments on the manuscript. We are grateful to Oliver Rocks for providing constructs, and Judith Trüb for help with cloning. We are grateful to the Microscopy Imaging Center of the University of Bern (https://www.mic.unibe.ch) and the Center for Microscopy and Image Analysis of the University of Zurich (https://www.zmb.uzh.ch) for support. This work was supported by the Swiss National Science Foundation Sinergia grant CRSII5_183550 to OP.

## Additional Information

The authors declare no competing interests.

