## Supplementary Figures for "Feedback regulation by the RhoA-specific GEF ARHGEF17 regulates actomyosin network disassembly"

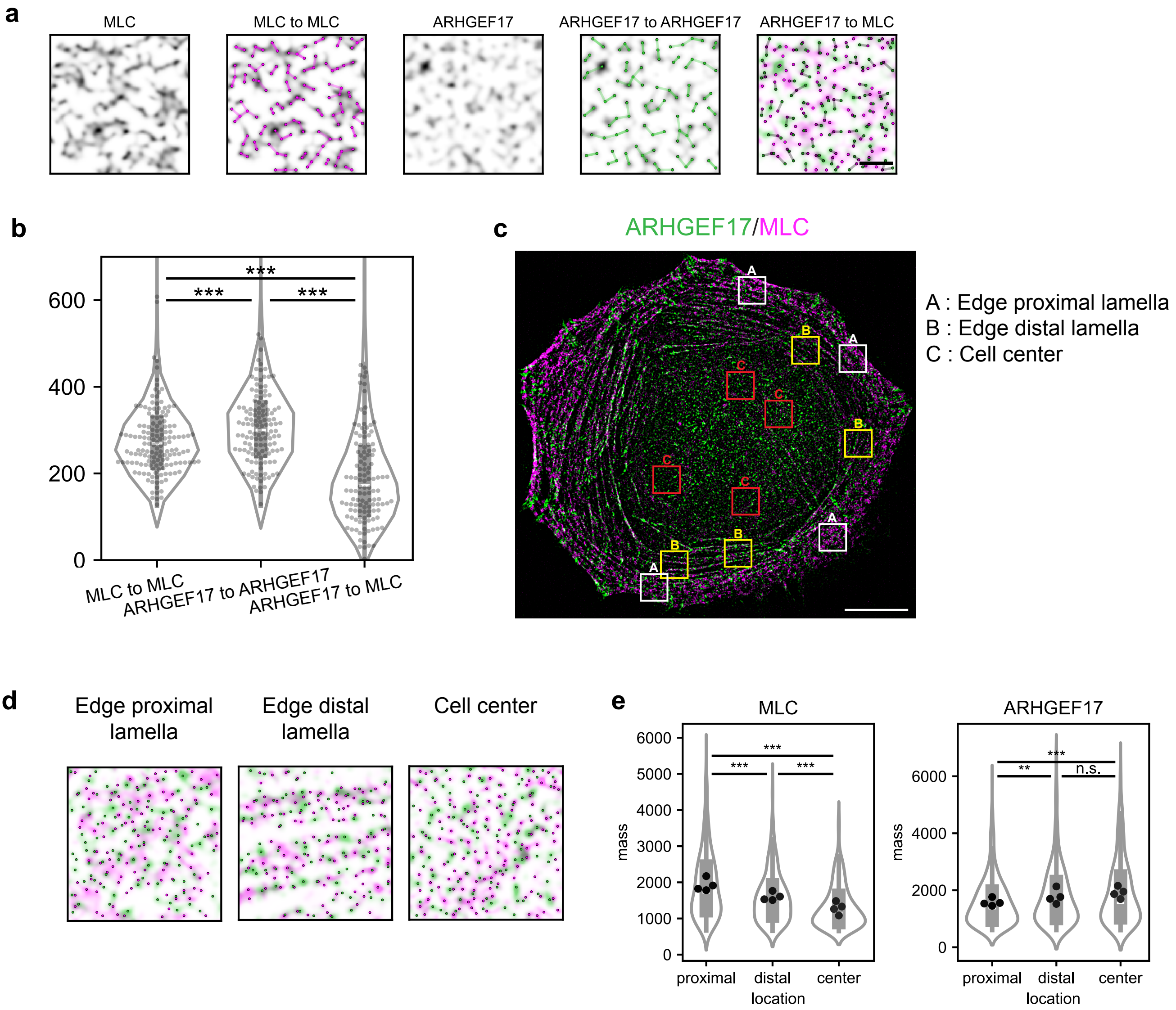


**Figure S1: Quantification of SIM MLC-mCherry/ARHGEF17-GFP images from Figure 2.**
**A:** Detection of MLC and ARHGEF17 spots, as gaussian blobs, and quantification of their distances to a nearest neighbour. A sample region of interest of both fluorescent signals and spot detection are shown. From left to right, raw channel MLC, MLC + spots + distances, raw channel ARHGEF17, ARHGEF17 + spots + distances, MLC and ARHGEF17 spots + distances.
**B:** Violin plots of the distribution of distances between directly neighbouring spots for MLC-MLC, ARHGEF17-ARHGEF17, and MLC-ARHGEF17.
C. Evaluation of MLC and ARHGEF17 spots in edge-proximal lamella, edge distal lamella, and cell center. ROIs for the measurements are shown over a MLC/ARHGEF17 overlay.
D. High magnification of ROIs in D with spot detection for specific ROIs: edge proximal and distal lamella, cell center.
E. Violin plots of distribution of the averaged spot fluorescence signals from 4 ROIs from (C)
Scale bar: (A) 2 µm,  (C) 10 µm

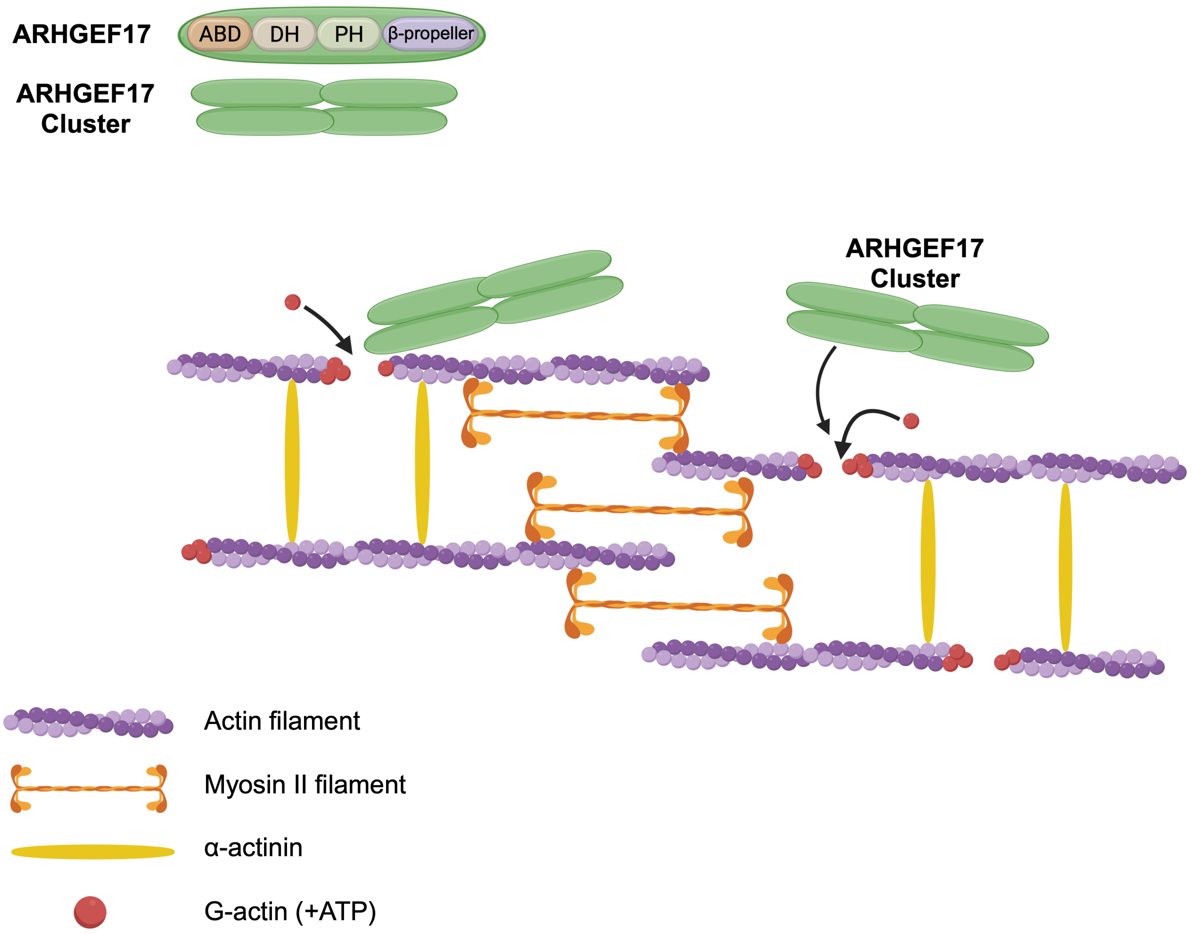


**Figure S2: Schematic depicting the organisation of ARHGEF17 clusters within the actomyosin lattice.** Actin filaments are crosslinked by myosin filaments. ARHGEF17 clusters bind the barbed end of actin filaments positioning it in between myosin heads (Figure 2, S1B). ARHGEF17 clusters are also mutually exclusive from alpha-actinin dimers that crosslink parallel actin filaments (Figure 3D-F).

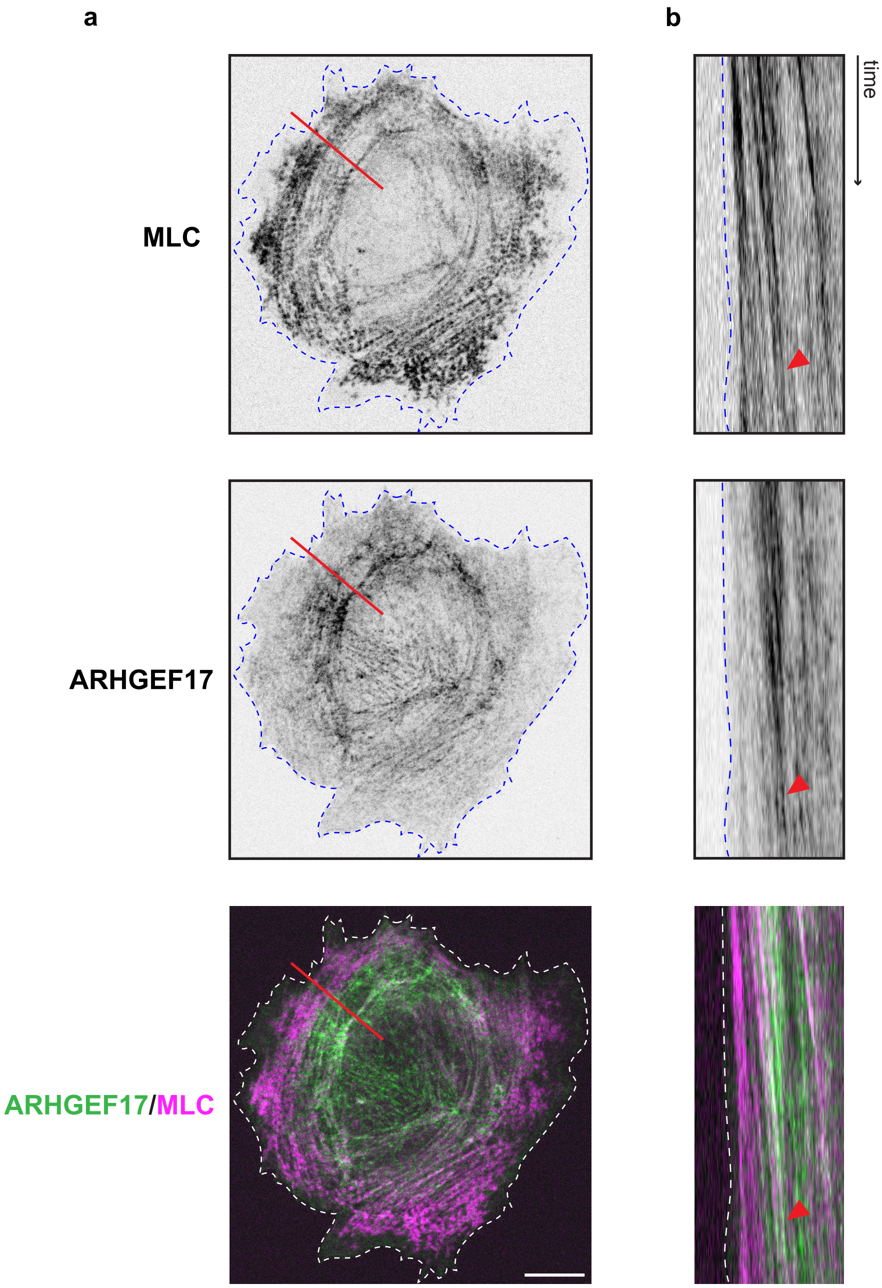


**Figure S3: Additional example of MLC/ARHGEF17 dynamics during lamella transverse arc disassembly.**
**A:** Spinning disk confocal timelapse imaging of a REF52 cell expressing ARHGEF17-GFP, and MLC-miRFP. Ibw contrast of single channels or color-coded overlays are shown.
**B:** Kymograph of line shown in A, red arrow depicts disassembly event.
Scale bar = 10 μm.

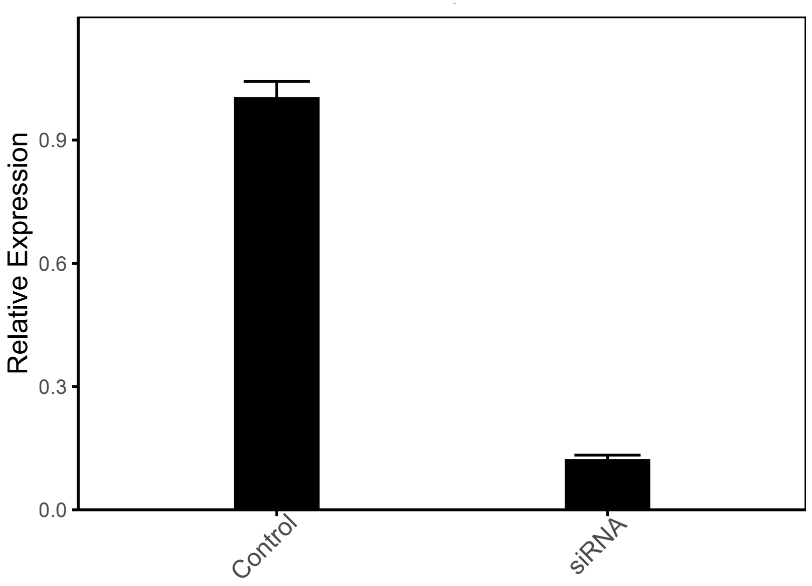


**Figure S4: Quantification of ARHGEF17 KD efficiency.**
Quantitative PCR (qPCR) analysis to evaluate the KD efficiency of ARHGEF17 KD. The expression levels of ARHGEF17 were determined using the ΔΔCt method and normalized to housekeeping gene GAPDH. The bar graph shows the mean relative expression for each condition, with the control set as the reference (relative expression = 1). Error bars represent the standard error of the mean (SEM) from two biological replicates.

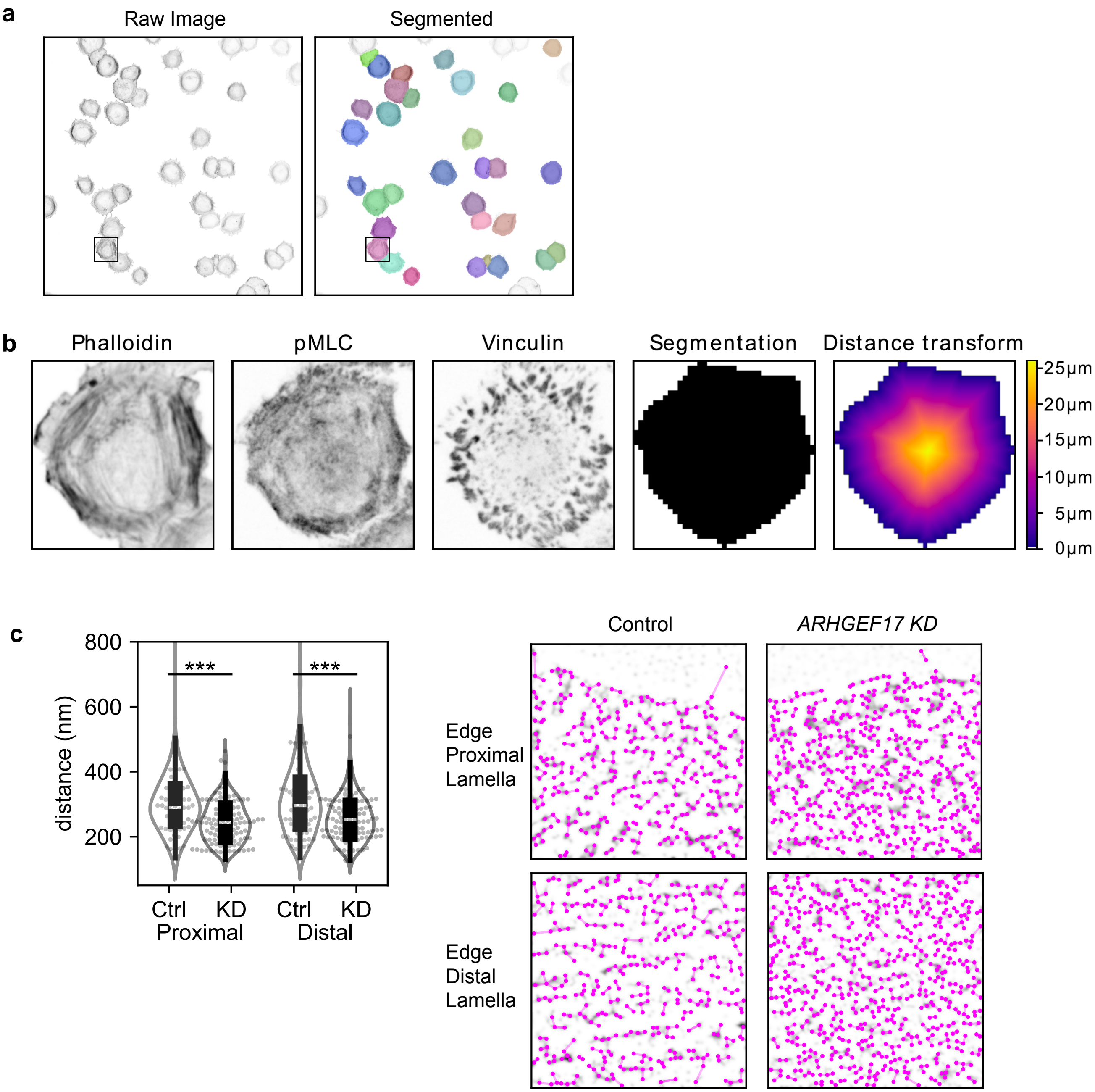


**Figure S5: Methodology for quantification of F-actin, pMLC and vinculin signals (relevant to Figure 5B) and MLC SIM data analysis in control versus ARHGEF17 KD cells**

**A:** Example field of view showing cell segmentation with cellpose in the phalloidin channel. Cells touching the image border are excluded from the analysis. The bounding box indicates the cell shown in panel B.

**B:** F-actin, vinculin and pMLC fluorescence profiles are extracted from single cells and population average fluorescence profiles using the distance transform are computed (see materials section).

**C:** Quantification of MLC distances in control versus ARHGEF17 KD cells using the identical method as in Figure S1. Violin plots of the distribution of distances between directly neighboring spots are shown.
